## Supplementary Material for "Fractal cycles of sleep: a new aperiodic activity-based definition of sleep cycles"

Yevgenia Rosenblum[1], Mahdad Jafarzadeh Esfahani[1], Nico Adelhöfer[1], Paul Zerr[1], Melanie Furrer[2], Reto Huber[2,3], Famke F. Roest[1], Axel Steiger[4], Marcel Zeising[5], Csenge G. Horváth[6], Bence Schneider[6], Róbert Bódizs[6], Martin Dresler[1]

[1] Radboud University Medical Centre, Donders Institute for Brain, Cognition and Behavior, Nijmegen, Netherlands,

[2] Child Development Center and Children's Research Center, University Children's Hospital Zürich, University of Zürich, Zürich, Switzerland,

[3] Department of Child and Adolescent Psychiatry and Psychotherapy, Psychiatric University Hospital Zurich, Zurich, Switzerland,

[4] Max Planck Institute of Psychiatry, Munich, Germany,

[5] Klinikum Ingolstadt, Centre of Mental Health, Ingolstadt, Germany,

[6] Semmelweis University, Institute of Behavioural Sciences, Budapest, Hungary

**Participants**

*Healthy adults*

We retrospectively analyzed polysomnographic recordings from the following studies (Table 1):

*Datasets 1 – 3:* 40, 40 and 33 healthy controls from three independent sleep studies in MDD conducted at the Max Planck Institute of Psychiatry, Germany. These datasets are described in Rosenblum et al. (2023 a) and Bovy et al. (2022). In addition, these participants are used as controls in MDD datasets A – C described below.

*Dataset 4:* 36 healthy participants from a home-based sleep study exploring simultaneous polysomnographic and EEG wearables conducted at the Donders Institute for Brain, Cognition and Behavior, the Netherlands (Described as Dataset 2 in Jafarzadeh Esfahani et al., 2023). The signal was recorded at participants' homes over three nights with a gap of a week between each recording. For consistency with other datasets (i.e., to end up with a comparable number of cycles provided by each participant), we used polysomnography (and not EEG recorded by wearables) from the first night only since it had the largest sample size (i.e., 5 subjects dropped out from the study after the first polysomnographic recording).

*Dataset 5:* 68 healthy controls from previous endocrinological studies conducted at the Max Planck Institute of Psychiatry, Germany, using only nights with no pharmacological or endocrine intervention. 60/68 participants are described in Rosenblum et al. (2023 b).

*Dataset 6:* 21 healthy children and adolescents from previous studies (Furrer et al., 2019; Volk et al., 2019; Jaramillo et al., 2020) conducted at the University Children's Hospital Zürich, Switzerland. For the control group to this dataset, we selected all healthy adults from Datasets 1 – 3, 5, 6 (n = 205) whose ages lay in the range of 23 – 25 years (the age when the brain maturation process is supposed to be finished (Giedd & Rapoport, 2010) and no age-related processes are expected to start). This resulted in 24 subjects with a mean age of 24.8 ± 0.9 years (Table S3 here).

In Rosenblum et al. (2023 a), Datasets A, B and C are referred to as the Replication Dataset 2, Main Dataset and Replication Dataset 1, respectively; in Bovy et al. (2022), the naming is the same as here.

*Patients with MDD*

We retrospectively analyzed polysomnographic recordings from our previous studies (Bovy et al., 2022; Rosenblum et al., 2023 a, Tables 1 – 2):

*Dataset A:* 40 long-term medicated MDD patients *vs.* 40 age- and gender-matched healthy controls (Dataset 1 here).

*Dataset B:* 38 MDD patients in unmedicated and 7-day medicated states *vs.* 40 healthy age and gender-matched controls (Dataset 2 here).

*Dataset C:* 33 MDD patients at 7-day and 28-day of medication treatment *vs.* 33 healthy age and gender-matched controls (Dataset 3 here).

Demographic and sleep characteristics of the patients, medication treatment and polysomnographic devices are described in our previous works (Bovy et al., 2022; Rosenblum et al., 2023 a). Here, Table S5 presents medication treatment. All studies were approved by the Ethics committee of the University of Munich. All patients gave written informed consent.

*Excluded participants*

Dataset 1: in one participant, the recording was available for the first 90 minutes only; in another participant, 25% of epochs were defined as "wake"; these two participants were excluded from the analysis. Dataset 2: in one participant, the recording was available for the first 133 minutes only; this subject was not included in the analysis. Dataset 3: in one participant 35% of all epochs were defined as "wake" and their data was excluded. Dataset 4: in one participant 50% of the epochs were tagged as "wake", another participant had no REM epochs, therefore, his classical sleep cycles could not be defined. These two participants were excluded. Dataset 5: in 5/68 participants, more than 30% of all epochs were tagged as “wake”. In addition, one participant had no REM epochs. These 6 participants were excluded from further analyses (Table 1). No pediatric and MDD participants were excluded.

Notably, in many sleep studies >25% of wake is not an exclusion criterion. However, here, we focus on sleep cycles specifically and can not assume that such a prolonged period of wakefulness during a night can be considered a part of a sleep cycle. An example of the data of one excluded participant is given in Fig.S6 C (S37).

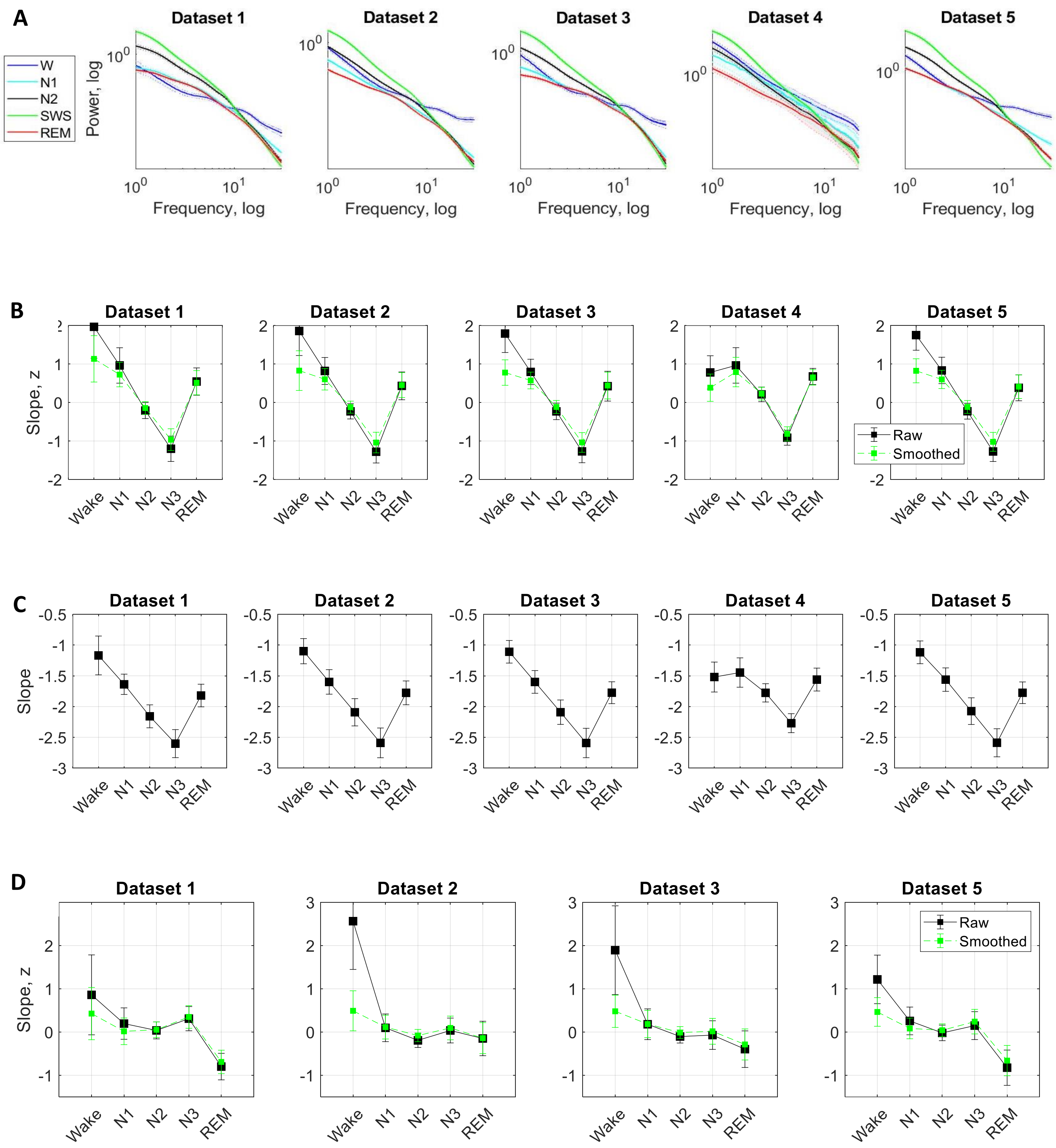
A
B
C
D
Dataset 1
Dataset 2
Dataset 3
Dataset 4
Dataset 5
W
N1
N2
SWS
REM
Power, log
Frequency, log
Slope, z
Slope
Raw
Smoothed
Wake
N1
N2
N3
REM

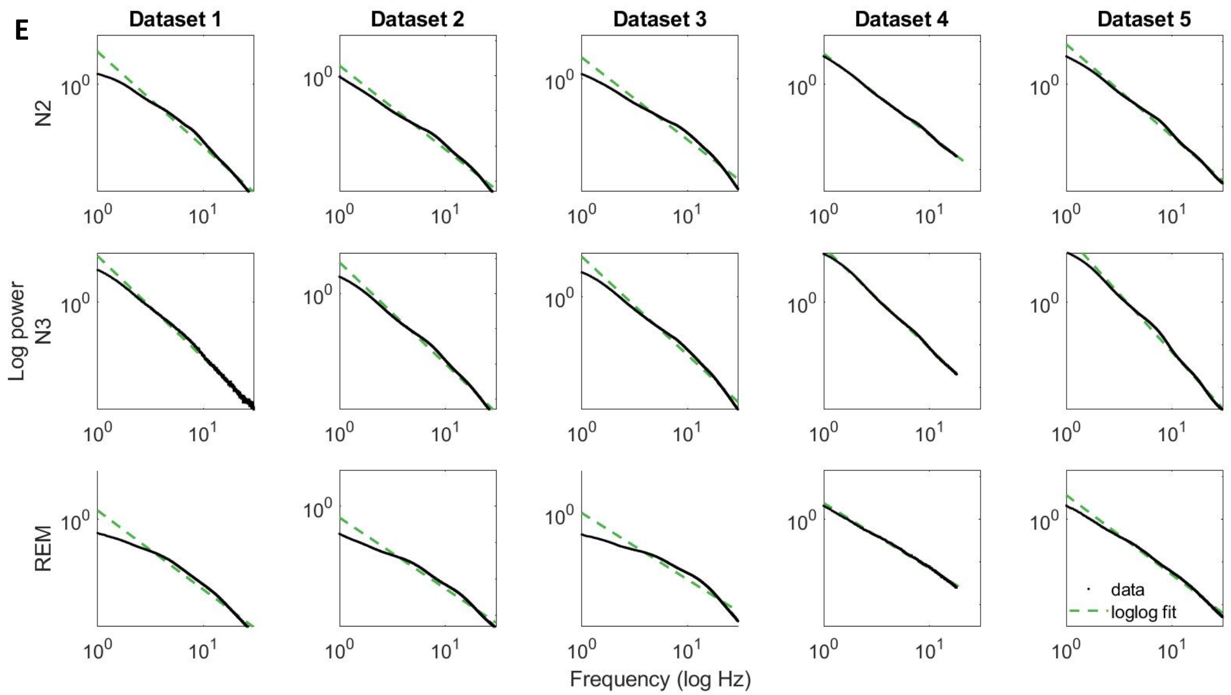

**Supplementary Figure 1. Stage-averaged fractal activity.**

**A: Fractal power component.** Fractal (aperiodic) power component averaged over each sleep stage separately over all participants of a given dataset is plotted as a function of frequency in the log-log space. Shading indicates standard errors. Slow-wave (green) and REM (red) sleep stages show the steepest and the flattest spectral decay, respectively in all datasets. For comparison, wake after sleep onset (blue) with the flattest decay in the higher frequency band is also shown. **B: 0.3 – 30Hz z-normalized slopes.** The slopes of the frontal fractal spectral power components in the 0.3 – 30Hz range are averaged over each sleep stage as defined by the hypnogram and z-normalized (black squares). For the analysis of the fractal cycles reported in the main text, fractal slopes were smoothened using the Savitzky-Golay filter (green squares). N3 is characterized by the steepest (most negative slopes) spectral decay compared to all other sleep stages in line with the existing literature. **C: 0.3 – 30Hz raw slopes.** Given that raw slope values contain valuable information per se (Bódizs et al., 2024), we also report raw slope values (before z-scoring). Note: Dataset 4 was filtered in the 0.3 – 18Hz range due to low-pass filtering during the recording. **D: 30 – 48Hz z-normalized slopes.** The slopes of the aperiodic (fractal) spectral power component in the 30 – 48Hz range are averaged over each sleep stage as defined by the hypnogram and z-normalized (black squares). Green squares show fractal slopes smoothened with the Savitzky-Golay filter. According to literature, REM sleep is expected to show the steepest (most negative) high-band slopes compared to all other sleep stages. However, we were able to replicate this finding in Datasets 1 and 5 only. Given poor differentiation between the stages in 2/4 datasets, this variable was not used in any analyses. Note: During recording, Dataset 4 was filtered in the 0.2 – 35Hz range and therefore, it was excluded from this subanalysis. **E: Log-log fit of data.** N – non-REM sleep, N3 – slow-wave sleep, REM – rapid eye movement sleep. Of note, in this study, we did not average spectral power/its slope over sleep stages; these graphs are shown to put our study in a broader context of the studies that looked at spectral power averaged over sleep stages for visualization only.

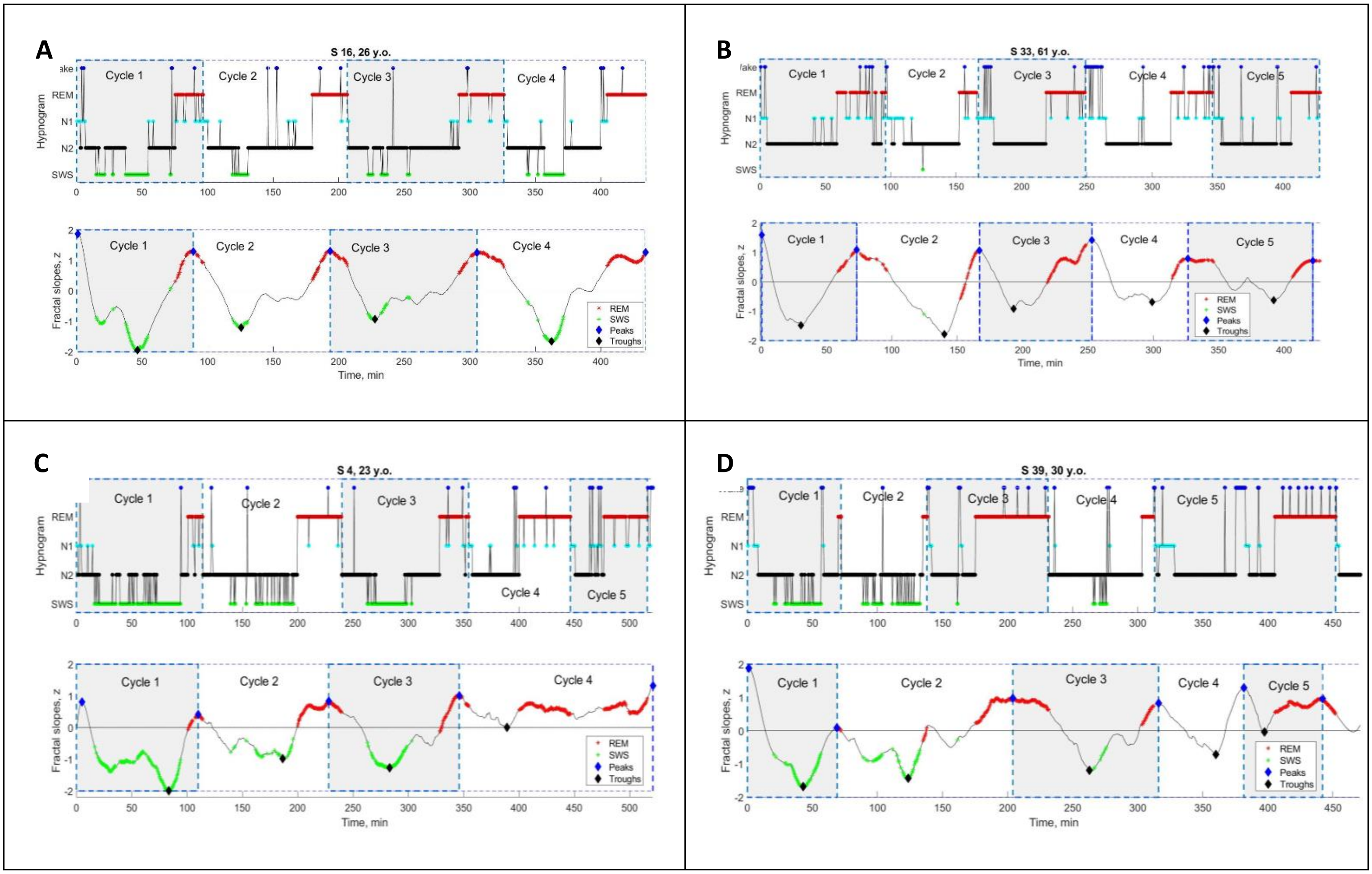

**Supplementary Figure 2. Individual fractal and classical sleep cycles in healthy adults.**

Time series of smoothed z-normalized fractal slopes (bottom) and corresponding hypnograms (top). The duration of the fractal cycle is an interval of time between two successive peaks (blue diamonds) defined with Matlab's function *findpeaks* with the minimum peak distance of 40 minutes and minimum peak prominence of 0.9 z. Two additional participants are presented in Fig.2 A – B. Supplementary PowerPoint File shared on https://osf.io/gxzyd presents these time series and hypnograms for all participants.

**A – B: one-to-one match.** In these two participants (from Dataset 3), there is an almost one-to-one match between fractal cycles defined by the algorithm and classical (non-REM – REM) cycles defined by the hypnogram.

**C – D: algorithm's misses.** Two participants from Dataset. In S4, the fourth fractal cycle corresponds to two classical cycles, No.4 and No.5, since the algorithm misses the local fractal peak at the 410th minute, which is not high enough (< |0.9| z). In S39, the second fractal cycle corresponds to two classical cycles, No.2 and No.3: the algorithm misses the local fractal peak at the 140th minute (the time of the corresponding REM episode), as the amplitude of the subsequent fractal descent is < |0.9| z. Two fractal cycles, No. 4 and No. 5, correspond to one classical cycle, No. 5: the algorithm identifies the wake episode at the 380th minute (in the middle of the 5th classical cycle) as a local fractal peak, i.e., the end of the fractal cycle. SWS – slow-wave sleep, REM – rapid eye movement sleep.

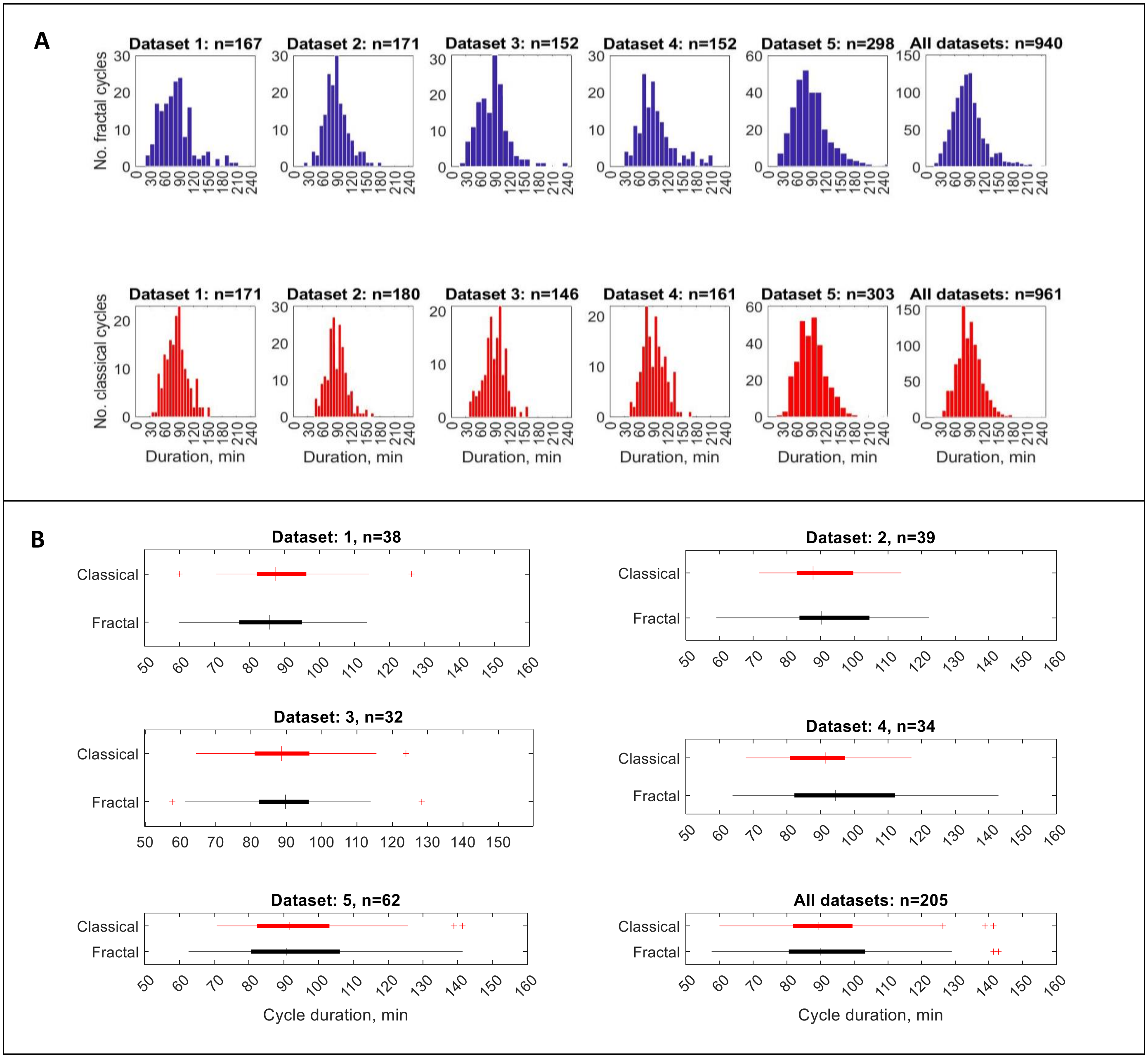

**Supplementary Figure 3. Group-averaged fractal and classical cycle durations.**

**A: Frequency distribution.** The individual fractal (top) and classical (bottom) cycles are counted (n) for each dataset separately and for all datasets merged. Across studies, 205 healthy adult participants provided 940 fractal cycles with a mean of 4.6 ± 1.0 cycles per participant and 961 classical cycles with a mean of 4.7 ± 0.9 cycles per participant. For both fractal and classical cycles, Kolmogorov-Smirnov test rejected the assumption that cycle duration comes from a standard normal distribution.

**B. Box plots.** In each box, a vertical central line represents the median, the left and right edges of the box indicate the 25th and 75th percentiles, respectively, the whiskers extend to the most extreme data points not considered outliers, and a plus sign represents outliers.

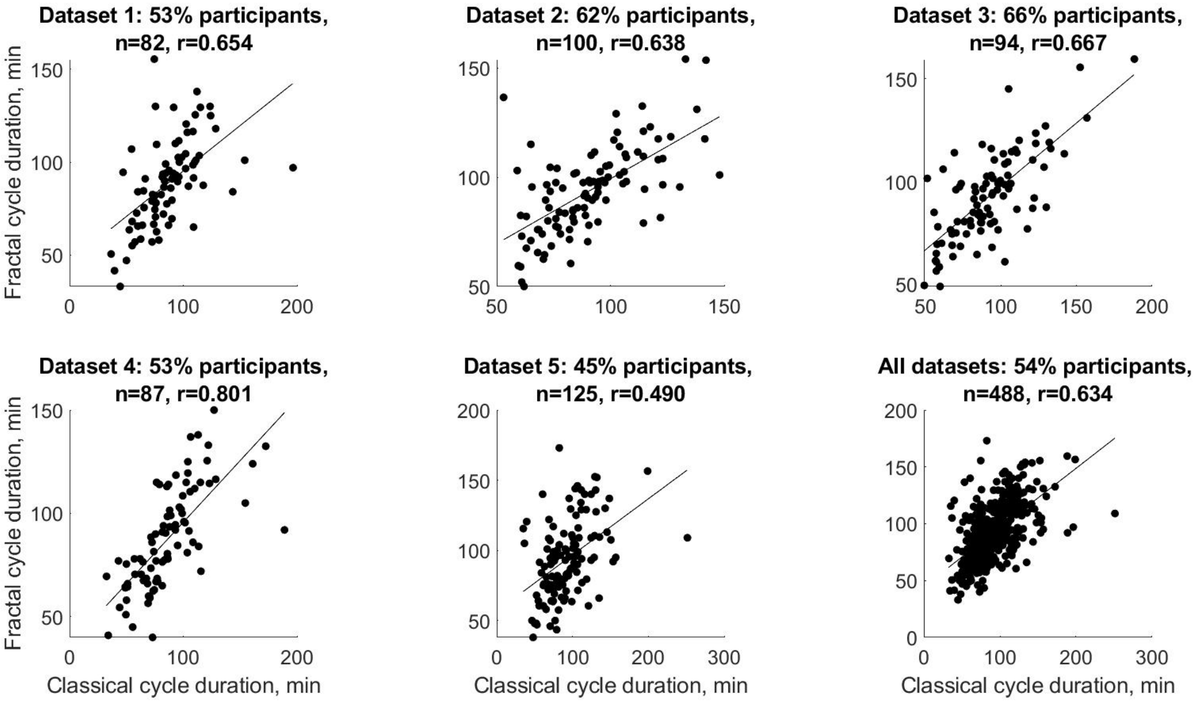

**Supplementary Figure 4. A subset of healthy adults with a one-to-one match between fractal and classical cycles durations** (correlations).

In a subset of the participants (from 45 to 66% in different datasets), there was a one-to-one match between fractal and classical cycles, each dot represents an individual cycle, n – number of cycles, all p-values < $10^{-9}$, r – Spearman's correlation coefficient.

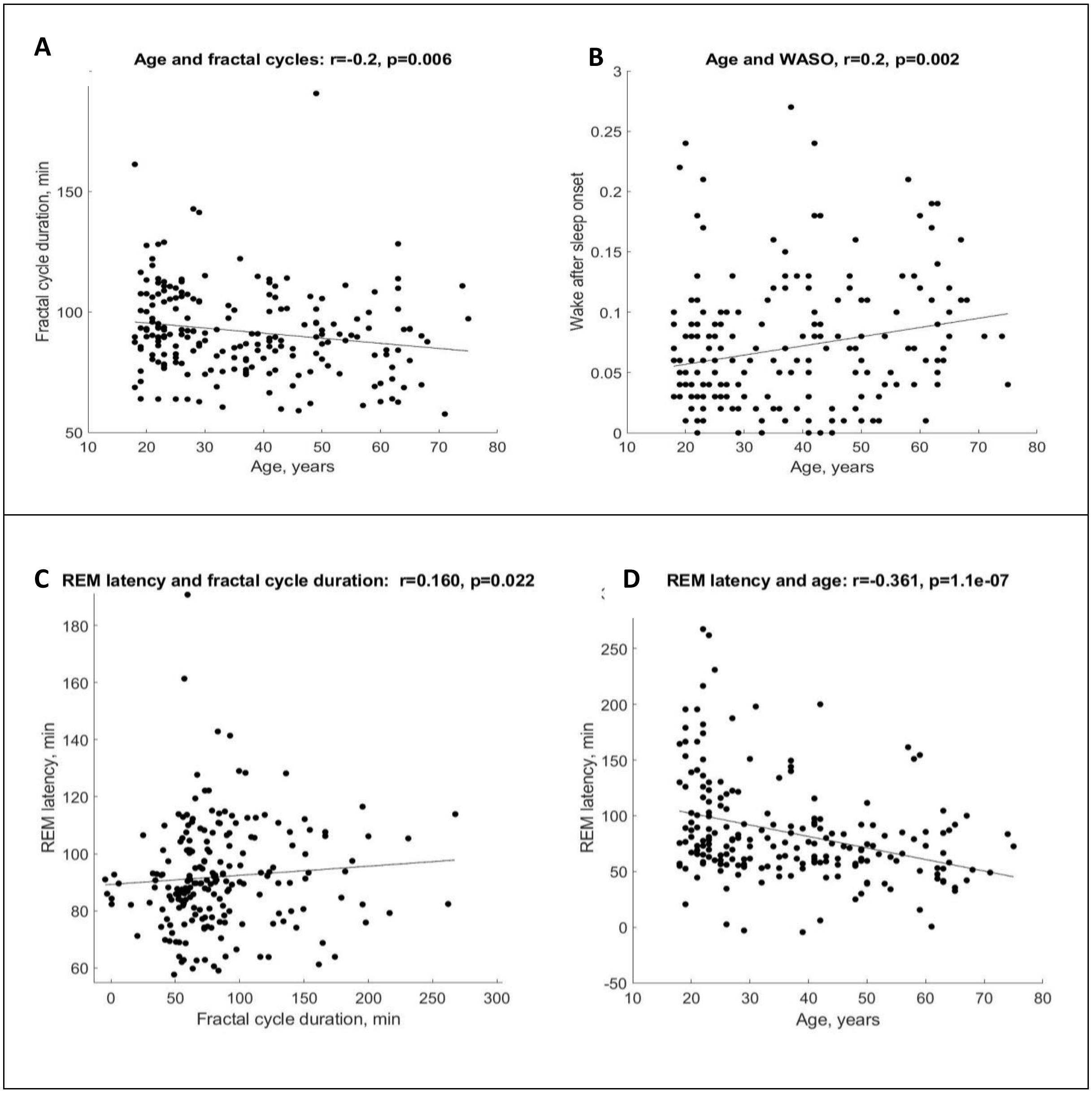

**Supplementary Figure 5. Correlations.**

**A:** Fractal cycle duration negatively correlates with the age of healthy participants.

**B:** The WASO proportion positively correlates with the age of healthy participants. The partial correlations between the fractal cycle duration and age adjusted for WASO and REM latency remained significant.

**C:** Fractal cycle duration positively correlates with REM latency of the healthy participants.

**D:** REM latency negatively correlates with the participant's age. The partial correlation between the fractal cycle duration and REM latency adjusted for the participant's age is non-significant. Age range: 18 – 75 years, median: 33.5 years, n = 205 (pooled Datasets 1 – 5), raw (non-ranked) values are presented, r – Spearman's correlation coefficient where values < 0.3 are considered as weak correlations, REM – rapid eye movement, WASO – wakefulness after sleep onset.

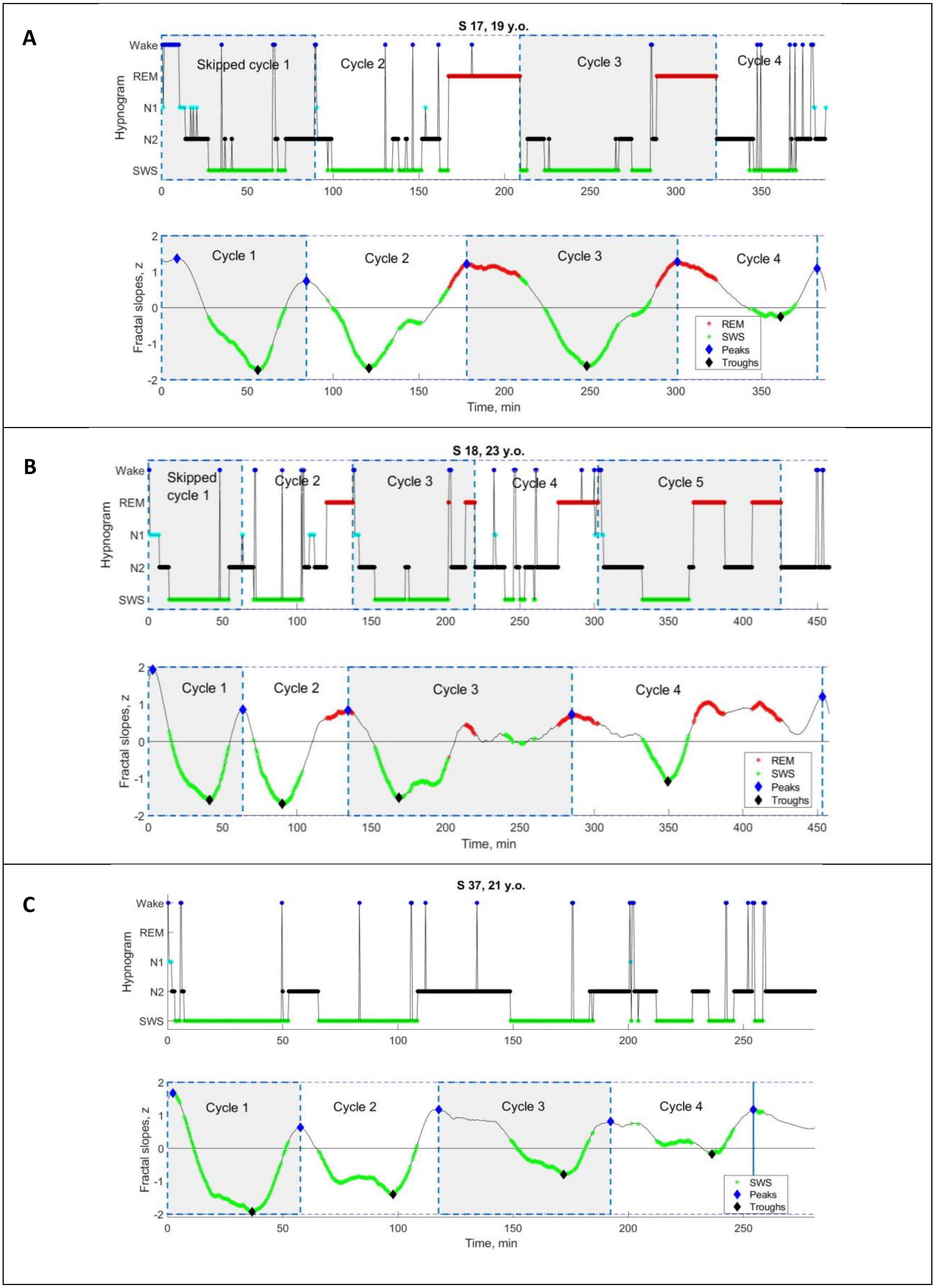
A
S 17, 19 y.o.
Skipped cycle 1
Cycle 2
Cycle 3
Cycle 4
Hypnogram
Wake
REM
N1
N2
SWS
Cycle 1
Cycle 2
Cycle 3
Cycle 4
Fractal slopes, z
REM
SWS
Peaks
Troughs
Time, min
B
S 18, 23 y.o.
Skipped cycle 1
Cycle 2
Cycle 3
Cycle 4
Cycle 5
Hypnogram
Cycle 1
Cycle 2
Cycle 3
Cycle 4
Fractal slopes, z
REM
SWS
Peaks
Troughs
Time, min
C
S 37, 21 y.o.
Hypnogram
Cycle 1
Cycle 2
Cycle 3
Cycle 4
Fractal slopes, z
SWS
Peaks
Troughs
Time, min

**Supplementary Figure 6. Individual cycles with skipped REM sleep.**

Time series of smoothed z-normalized fractal slopes (bottom) and corresponding hypnograms (top) observed in three young healthy participants (from Dataset 4). Hypnograms show skipped first cycles (as well as the rest of the classical cycles).

**A:** In S17, at the 90th minute, an episode of REM sleep is expected to appear – except only a "lightening of sleep" (wake, N1 and N2) is observed. We divided the long 209-minute cycle into two cycles, the 90-minute skipped cycle and the 119-minute normal cycle.

**B:** In S18, at the 63rd minute, an episode of REM sleep is expected to appear – except only a "lightening of sleep" (N1 and N2) is observed. We divided the long 138-minute cycle into two cycles, the 63-minute skipped cycle and the 75-minute normal cycle. The fractal cycle algorithm was very effective in detecting skipped cycles, showing a one-to-one match with divided – but not long undivided – cycles.

**C:** S37's hypnogram shows that she has no REM sleep at all, i.e., all her cycles are the "skipped" ones. Based on this, S37 was even excluded from the formal analysis. This example is presented here to illustrate that the fractal cycle algorithm is sensitive enough in detecting sleep cycles even in the absence of REM sleep.

SWS – slow-wave sleep, REM – rapid eye movement sleep.

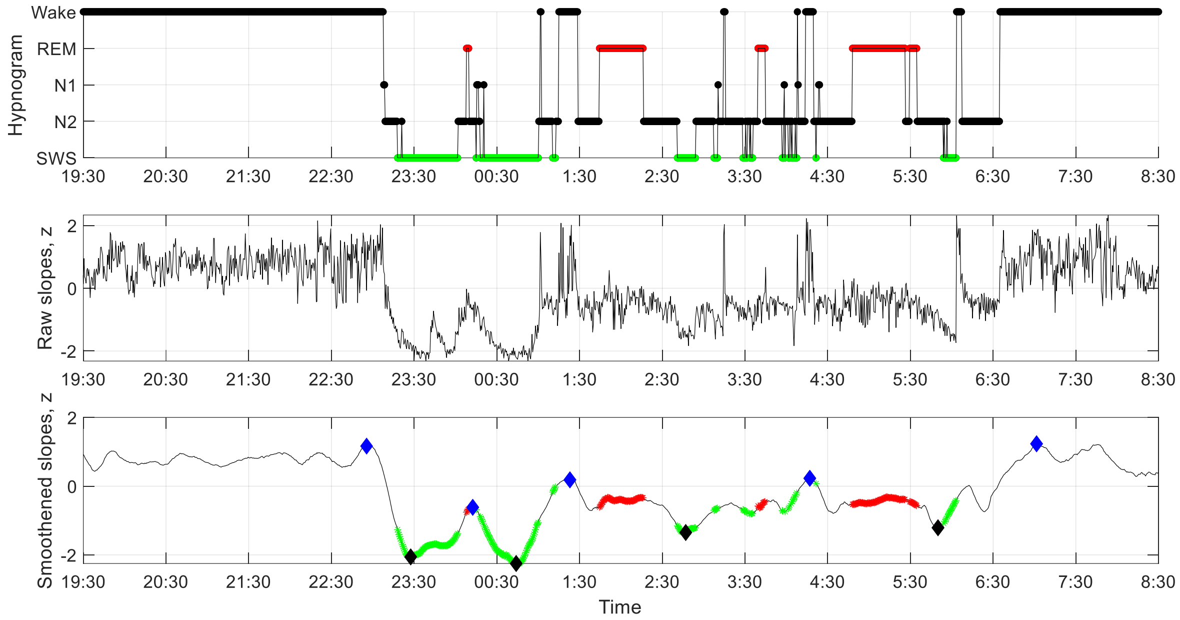

**Supplementary Figure 7. Individual fractal time series across 13 hours.**

Time series of raw (middle) and smoothed z-normalized fractal slopes (bottom) as well as the corresponding hypnograms (top) in a 25-year-old healthy male. In addition to sleep-related fractal activity, this figure shows fractal activity during 3 hours before the sleep onset and 2 hours after awakening. The graph shows that fractal cycles are not observed during wake, being specific to sleep. This participant does not come from the datasets depicted in the current study (where no recordings > 8h were available). The study, EEG device and preprocessing are described in Rosenblum et al., 2024. EEG power was averaged over F4, C4, and O2 electrodes, differentiated into its components, z-scored and smoothed as described in Methods of the current paper. The duration of the fractal cycle is a time interval between two successive peaks (blue diamonds). SWS – slow-wave sleep (green dots), REM – rapid eye movement sleep (red dots).

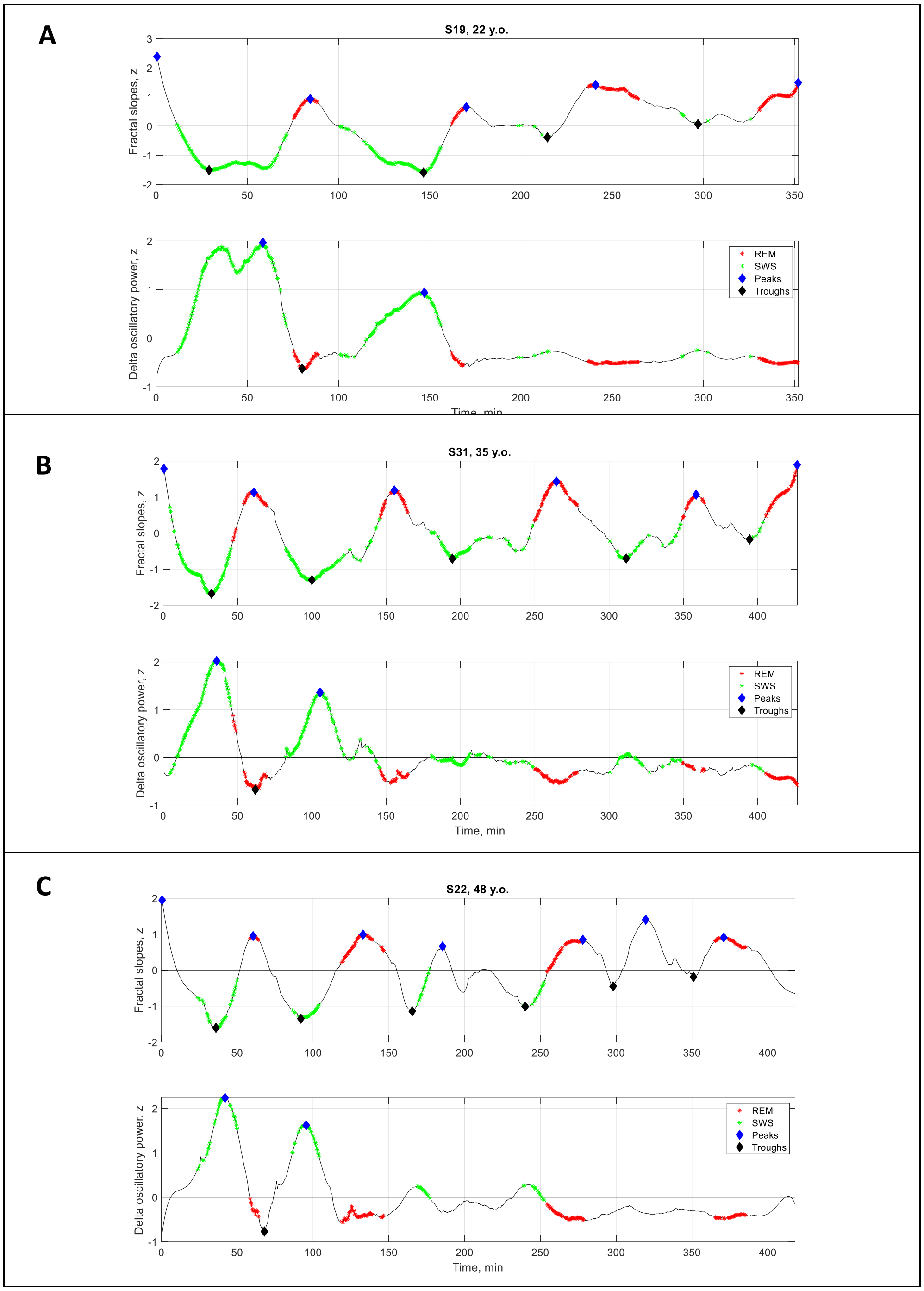

**Supplementary Figure 8. Individual fractal *vs.* SWA time series.**

Time series of smoothed z-normalized fractal slopes (top) and SWA/delta (1 – 4 Hz) oscillatory power (bottom) observed in three healthy adults (from Dataset 3). SWS – slow-wave sleep, SWA – slow-wave activity, REM – rapid eye movement sleep.

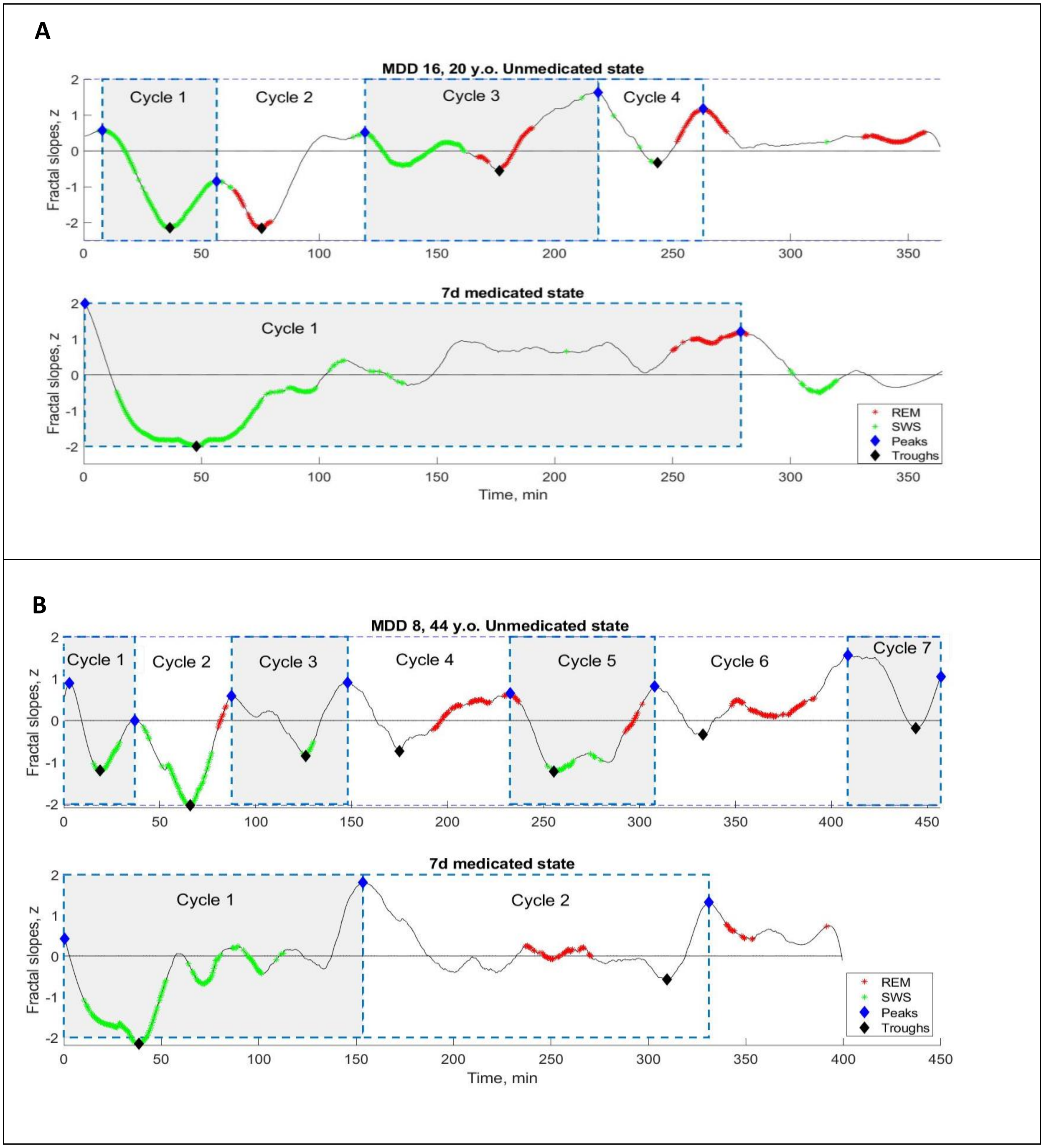

**Supplementary Figure 9. Individual fractal cycles in MDD patients.**

Time series of smoothed z-normalized fractal slopes observed in two additional MDD patients (Dataset B, an additional patient is presented in Fig.4A) in their unmedicated (top) and 7-day medicated (bottom) states. Fractal cycles duration (defined as an interval of time between two successive peaks, blue diamonds) is longer in the medicated compared to unmedicated states, reflecting shallower fluctuations of fractal (aperiodic) activity. MDD – major depressive disorder, SWS – slow-wave sleep, REM – rapid eye movement sleep.

**Fractal cycles in patients with insomnia: a pilot**

We compared fractal cycle duration in 11 patients with insomnia (18.18% male; age: 44 ± 13.2 years, n = 11, No. cycles = 51) and 11 healthy controls (54.5% male; age: 42.4 ± 15.4 years, n = 11, No. cycles = 46), using open access dataset from a cross-sectional study on psychophysiological insomnia (Rezaei et al., 2017). The analysis were performed as described in Methods. However, due to low-pass filtered EEG data, the fitting of spectral slopes was performed in the 1 – 18Hz range.

An individual example of smoothed fractal slope time series and hypnograms is shown in Fig.S10 A. We found that patients with insomnia showed a shorter duration of fractal cycles compared to controls with a medium effect size (83 ± 45 *vs.* 101 ± 43 min, p = 0.04, Cohen's d = -0.4, 4.6 cycles/participant *vs.* 4.2 cycles/participant, Fig.S10 B – C).

These findings are in line with the existing literature on flatter slopes in insomnia patients compared to controls (Andrillon et al., 2020 and Fig. S10 D), strengthening the hyperarousal model of insomnia.

This analysis is an outlook only and future studies using a higher sample size should confirm this finding.

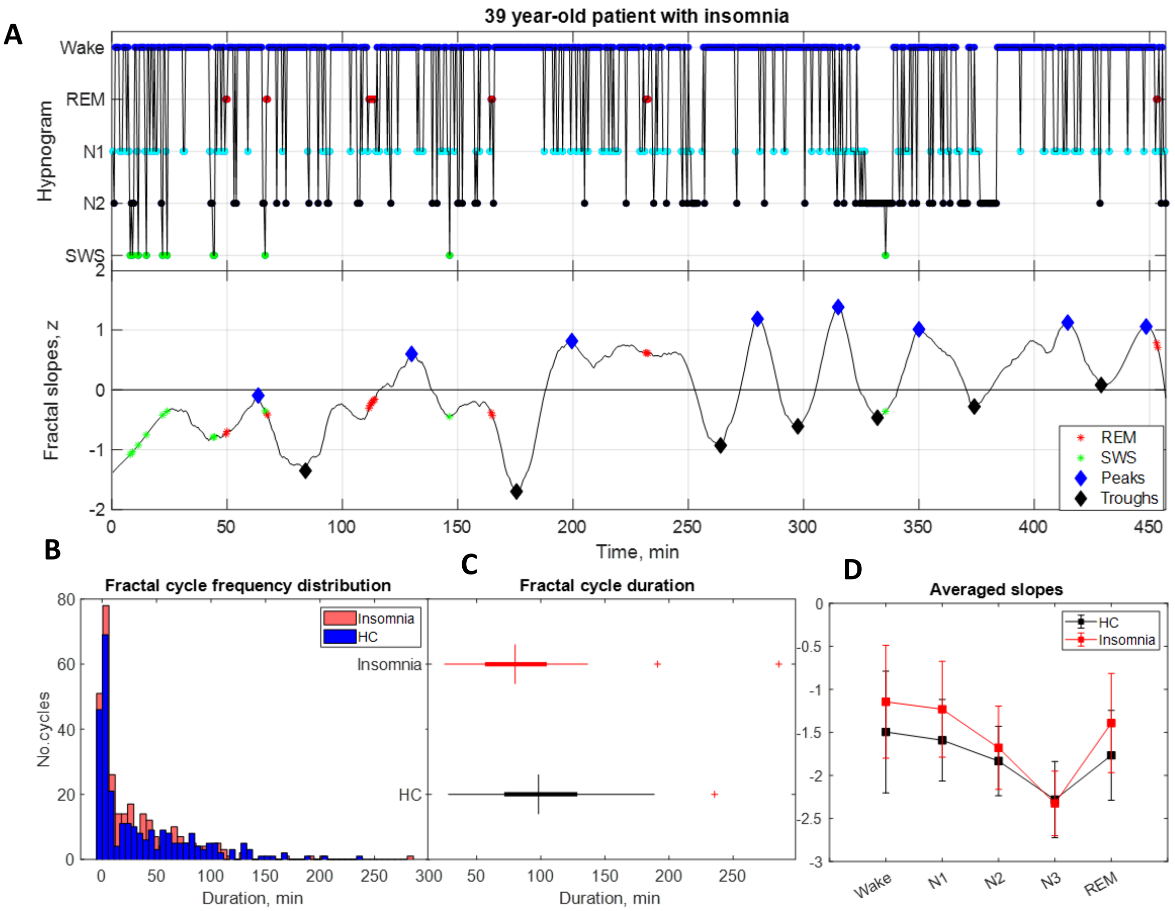

**Supplementary Figure 10. Fractal cycles in patients with insomnia.**

**A. Individual example:** time series of smoothed z-normalized fractal slopes (bottom) and corresponding hypnograms (top). The duration of the fractal cycle is a time interval between two successive peaks (blue diamonds) defined with Matlab's function *findpeaks* with a minimum peak distance of 20 minutes and minimum peak prominence of 0.9 z. SWS – slow-wave sleep, REM – rapid eye movement sleep. **B. Histograms**: The frequency distribution of fractal cycle durations in patients with insomnia compared to controls. Kolmogorov-Smirnov's test rejected the assumption that cycle duration comes from a standard normal distribution. **C. Box plots:** in each box, a vertical central line represents the median, the left and right edges of the box indicate the 25th and 75th percentiles, respectively, the whiskers extend to the most extreme data points not considered outliers, and a plus sign represents outliers. Patients with insomnia show shorter fractal cycle duration compared to controls**. D. Averaged slopes:** The slopes of the frontal fractal spectral power components in the 1 – 18Hz range are averaged over each sleep stage as defined by the hypnogram. N3 is characterized by the steepest (most negative slopes) spectral decay compared to all other sleep stages in line with the existing literature. Patients present with flatter slopes compared to controls in all sleep stages confirming previous findings (Andrillon et al., 2020).

**Table S1: Fractal cycle topography in healthy adults**

| Area | Dataset 1 | Dataset 2 | Dataset 3 | Dataset 4 | Dataset 5 |
|---|---|---|---|---|---|
| **Classical sleep cycle duration, min** | | | | | |
| --- | 86.2 ± 23.3 | 90.0 ± 21.3 | 89.0 ± 22.7 | 92.2 ± 23.7 | 91.9 ± 29.0 |
| **Fractal sleep cycle duration, min** | | | | | |
| F | NA | 90.0 ± 25.5 | 86.4 ± 31.2 | 94.7 ± 37.1 | 89.9 ± 37.1 |
| C | 86.4 ± 35.2 | 91.1 ± 29.4 | 85.2 ± 34.2 | 95.4 ± 37.3 | 90.8 ± 39.9 |
| P | NA | 95.0 ± 32.3 | 89.8 ± 37.2 | NA | 90.7 ± 41.9 |
| O | NA | 89.7 ± 28.6 | 85.0 ± 31.3 | 100.0 ± 47.0 | 92.2 ± 42.5 |
| **Classical-fractal cycles correlation, r coefficient** | | | | | |
| F | NA | 0.508 | 0.565 | 0.474 | 0.513 |
| C | 0.331 | 0.364 | 0.213 | 0.478 | 0.277 |
| P | NA | 0.273 | 0.120 | NA | 0.411 |
| O | NA | 0.306 | 0.239 | 0.516 | 0.279 |
| **Classical-fractal cycles correlation, p-value** | | | | | |
| F | NA | 0.001 | 0.001 | 0.005 | <0.001 |
| C | 0.042 | 0.023 | 0.242 | 0.004 | 0.029 |
| P | NA | 0.093 | 0.512 | NA | <0.001 |
| O | NA | 0.058 | 0.189 | 0.002 | 0.028 |

*F – frontal (averaged over F3 and F4), C – central (averaged over C3 and C4), P – parietal (averaged over P3 and P4), O – occipital (averaged over O1 and O2) electrodes.*

**Table S2: Fractal cycle characteristics: frequency bands comparison**

| **Parameter** | **0.3 – 30Hz** | **1 – 30Hz** |
|---|---|---|
| Fractal cycles, No./night | 4.6 ± 1.0 | 4.6 ± 1.1 |
| Fractal sleep cycle duration, min | 89.1 ± 34.0 | 90.7 ± 36.9 |
| Descent amplitude, z | -2.2 ± 0.8 | -2.1 ± 0.8 |
| Ascent amplitude, z | 2.2 ± 0.6 | 2.1 ± 0.6 |
| Classical – fractal cycles duration correlation, r | 0.488 | 0.397 |
| Classical – fractal cycles duration correlation, p | 10e-13 | 10e-9 |

*Mean values ± SD (min) are presented for the pooled dataset (n = 205). The 1 – 30Hz band is added to control for a possible distortion (the so called "knees'' of the spectrum) of the linear fit by excluding low frequencies with strong oscillatory activity. Both bands, however, show similar results probably because of the smoothening procedure used in this study.*

**Table S3: Pediatric demographic and sleep characteristics**

| Characteristic | Children and adolescents (Dataset 6) | Controls: young adults (from Datasets 2, 4, 5) |
|---|---|---|
| Sample size | 21 | 24 |
| Age, years | 12.4 ± 3.1 | 24.8 ± 0.9 |
| Age range, years | 8 – 17 | 23 – 25 |
| Wake, % | 4.94 | 6.09 |
| Non-REM stage 1, % | 3.29 * | 7.27 |
| Non-REM stage 2, % | 41.89 | 42.23 |
| Slow-wave sleep, % | 31.06 * | 24.92 |
| REM sleep, % | 18.82 | 18.58 |
| Total sleep time, min | 444 ± 37 | 441 ± 44 |
| Classical sleep cycle duration, min | 80.4 ± 23.0 * | 89.8 ± 22.2 |
| Fractal sleep cycle duration, min | 75.5 ± 33.7 * | 94.1 ± 32.1 |
| Descent amplitude, z | -2.3 ± 0.9 | -2.2 ± 0.8 |
| Ascent amplitude, z | 2.2 ± 0.7 | 2.1 ± 0.6 |
| No. fractal cycles | 112 | 121 |
| No. classical cycles | 112 | 114 |

*± shows mean and SD, * indicates statistically significant differences as revealed by the non-parametric test, REM – rapid eye movement.*

**Autocorrelation and partial autocorrelation**

*Method*

To further explore the fluctuating nature of the fractal slope time series, we assessed the autocorrelation and partial autocorrelation patterns of this data using Matlab's *autocorr* and *parcorr* functions, respectively. In autocorrelation, a given value from a time series is regressed on previous values from that same time series. Partial autocorrelation is similar to autocorrelation except that it displays only the correlation between two observations that the shorter lags between those observations do not explain. In other words, the partial correlation for each lag is the unique correlation between those two observations after partialling out the intervening correlations, i.e., it controls for other lags.

For both autocorrelation and partial autocorrelations, we defined the number of lags as 180, which corresponds to 90 minutes, an average duration of a fractal cycle, with a 30-second step. We assessed these correlations in each participant separately and then averaged the correlation coefficients for each time lag over all participants of a given dataset and in a pooled dataset.

*Results*

Autocorrelation strength decayed throughout the lags showing a somewhat sinusoid shape. Specifically, positive correlations of moderate strength were observed for the 0.5 – 14 minute lags; for the 14 – 25 minute lags, the correlations were weak. In addition, weak positive autocorrelations were observed around the 90th minute while weak negative autocorrelations were observed around the 45th minute (Fig. S11, left), corroborating 90-minute periodicity of fractal cycles.

The partial autocorrelation that controls for other lags further revealed that only 0 – 5 minute lag coefficients were statistically significant, i.e., autocorrelation equals zero at lags greater than 5 minutes (Fig. S11, right). This finding indicates that the fractal slopes of the consecutive sleep epochs within a given 5 minutes are not independent, they autocorrelate so that they are much more likely to appear in an observed pattern (e.g., as in Fig.2 A) than expected by chance.

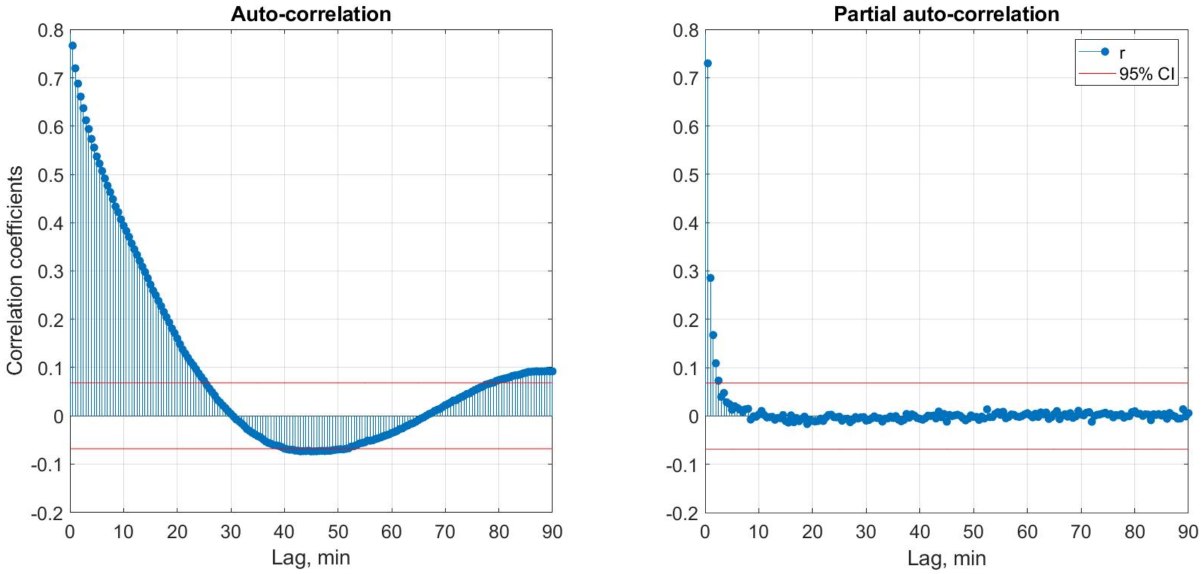

**Supplementary Figure 11.** Autocorrelation **(left)** and partial autocorrelation **(right)** of the time series of the fractal slopes averaged over each 30 seconds of sleep show the correlation of this time series with a delayed version of itself as a function of time lag. Red horizontal lines indicate approximate 95% probability limits for a purely random process. Correlation coefficients were calculated separately for each participant and then averaged over the pooled dataset (n = 205).

**Cross-correlations**

*Method*

To further model the temporal relationship between fractal and classical cycles, we explored them on a finer grained level using cross-correlations between time series of fractal slopes and time series of non-REM or REM sleep proportion per each 5 minutes of sleep. Cross-correlation allows one to evaluate how two time series might concomitantly covary in the same or opposite directions at given temporal intervals (i.e., lags). Thus, it enables the identification of temporally coordinated fluctuations between two variables. From the shape of the cross-correlation function, information concerning the possible direction of the influence between the two processes can be obtained.

Cross-correlation analyses were performed between the time series of the fractal slopes averaged over each 5 minutes (based on the results of autocorrelation analysis reported above) of sleep on one side and the time series of the proportion of either REM or non-REM stages 2 and 3 (together) on the other side. We averaged the values of all the time series (originally calculated for each 30 seconds of sleep) over each 5 minutes of sleep. The cross-correlation coefficients were computed for lags ranging from -40 to 40 minutes, a period approximately corresponding to an average duration of a fractal cycle, with a 5-minute step. Confidence intervals of 95% were calculated to infer statistical significance. We calculated cross-correlations in each participant separately and then averaged correlation coefficients for each time lag over all healthy adult participants in a pooled dataset (Datasets 1 – 5). Likewise, we calculated the proportion of the participants who showed statistically significant correlation coefficients for each lag.

*Results*

We found that the time series of fractal slopes positively cross-correlated with the time series of REM sleep proportion and negatively correlated with the time series of non-REM sleep proportion for lags lying between -15 and +5 minutes (Fig. S12, Table S4). At the individual level, significant correlations were observed in more than 80% of the participants for the -10 – 0-minute lags. Bayesian prevalence analysis revealed that the maximum *a posterior* prevalence estimate is equal to 0.79 while the Bayesian highest posterior density interval (true population level) with 96% probability level lies within the 0.73 – 0.85 range.

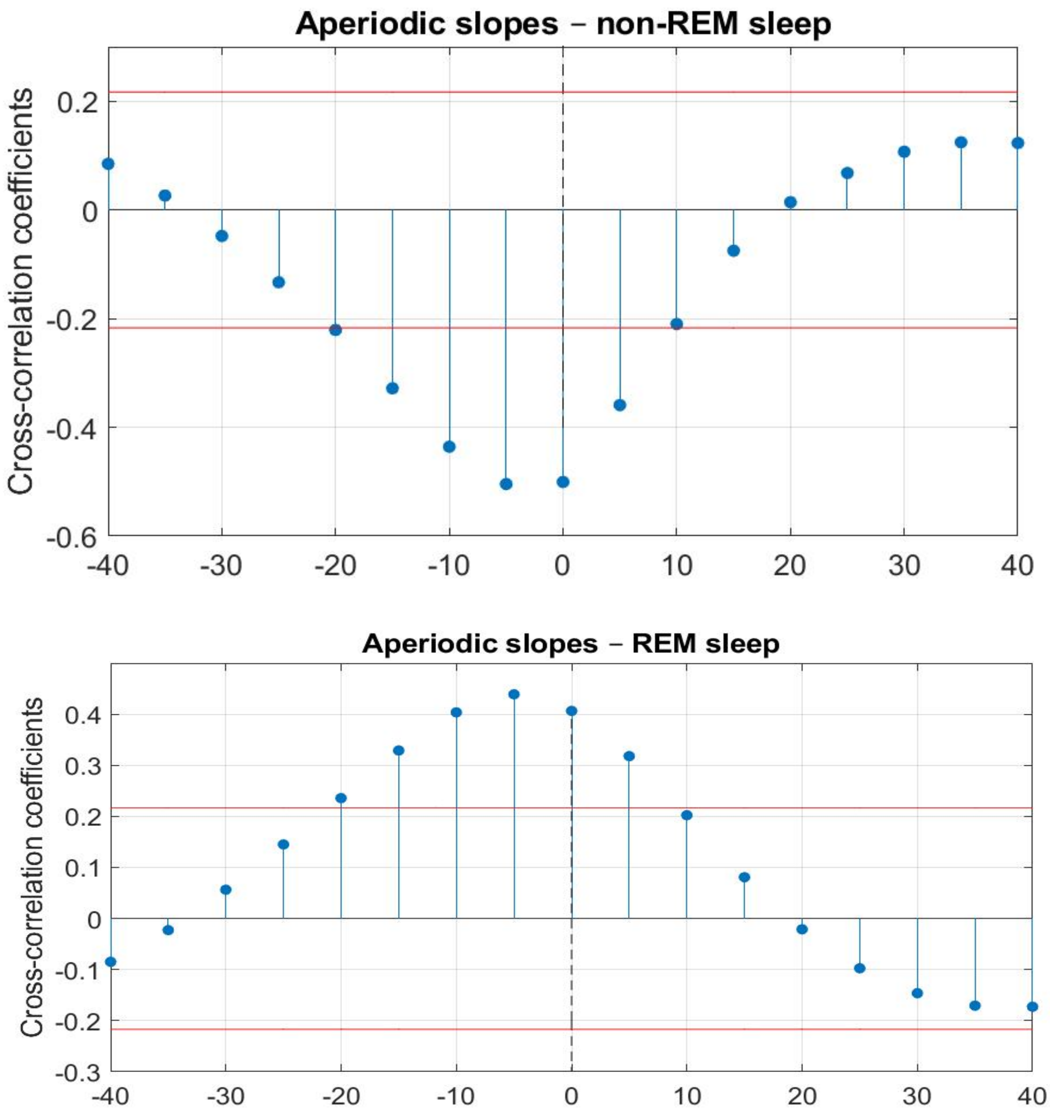

**Supplementary Figure 12. Cross-correlations.** Cross-correlations between the time series of fractal slopes on the one side and the proportion of non-REM **(top)** or REM sleep **(bottom)** per 5 minutes on the other side. Cross-correlations were calculated for each participant individually, then correlation coefficients were averaged over all healthy adults (n = 205). Negative and positive lags mean that fractal slope time series are leading and lagging, respectively. Here, the shape of the cross-correlation function does not allow one to decide which time series is leading and which one is lagging here. The horizontal red lines mark the CI of 95%, the absolute values above these lines indicate statistical significance (r > |0.22|). REM – rapid eye movement sleep, CI – confidence interval.

**Table S4: Cross-correlations**

| Time lag | non-REM<-Slopes | Slopes<-non-REM | REM<-Slopes | Slopes<-REM |
|---|---|---|---|---|
| Correlation coefficients, r | | | | |
| 0 min | -0.50 | | 0.40 | |
| 5 min | -0.50 | -0.36 | 0.44 | 0.32 |
| 10 min | -0.44 | n.s. | 0.40 | n.s. |
| 15 min | -0.33 | n.s. | 0.33 | n.s. |
| 20 min | n.s. | n.s. | n.s. | n.s. |
| Participants showing significant effect, % | | | | |
| 0 min | 85 | | 80 | |
| 5 min | 85 | 70 | 87 | 72 |
| 10 min | 84 | 54 | 86 | 53 |
| 15 min | 76 | 21 | 79 | 18 |
| 20 min | 52 | 9 | 5 | 61 |

*REM – rapid eye movement sleep, n.s. – non-significant. Only lags associated with statistically significant correlation coefficients are reported, r's higher than 0.7 are considered as strong correlation scores, values lower than 0.3 are considered as weak, r's values in the range of 0.3 – 0.7 are considered as moderate scores. Correlation coefficients were calculated for each healthy adult individually and then averaged over all participants.*

**Table S5: Demographic and clinical characteristics of the subgroups of patients by medication class (mean ± SD)**

| Medication class | n | Age | No. females | No. of previous depressive episodes | HAM-D baseline | HAM-D 7-day |
|---|---|---|---|---|---|---|
| SSRI (citalopram, escitalopram, paroxetine, sertraline) | 13 | 29.9±10.0 | 8 | 0.6±0.8 | 19.7±4.2 | 13.9±4.6 |
| TCA (trimipramine, amitriptyline, amitriptylinoxide) | 8 | 36.6±11.9 | 4 | 2.1±1.1 | 22.1±3.4 | 16.6±5.5 |
| NDRI (bupropion) | 6 | 30.7±10.5 | 3 | 0.7±0.5 | 18.5±3. 5 | 17.8±3.2 |
| SNRI (venlafaxine, duloxetine) | 6 | 31.7±10.9 | 2 | 1.7±0.83 | 18.3±2.5 | 14.7±5.5 |
| NaSSA (mirtazapine) | 5 | 26.8±6.1 | 3 | 2.6±3.2 | 20.2±4.8 | 13.8±4.7 |
| REM suppressive (SSRI, SNRI, amitriptyline, amitriptylinoxide) | 21 | 31.1±10.3 | 11 | 1.1±1.0 | 19.2±3.6 | 14.1±4.5 |
| REM non-suppressive (trimipramine, bupropion, mirtazapine) | 17 | 31.6±10.4 | 7 | 1.7±2.0 | 20.7±4.1 | 16.6±4.9 |

*HAM-D – Hamilton Depression Rating Scale, REM – rapid eye movement sleep, SD – standard deviation, NaSSA – noradrenergic and specific serotonergic antidepressants, NDRI – norepinephrine-dopamine reuptake inhibitor, SNRI – serotonin-norepinephrine reuptake inhibitors, SSRI – selective serotonin reuptake inhibitors, TCA – tricyclic antidepressants.*

**Table S6: Fractal cycle topography in MDD**

| Data set | Group | F | C | P | O |
|---|---|---|---|---|---|
| A | Healthy controls | NA | 84 ± 35 | NA | NA |
| | long-termed med. MDD | NA | **97 ± 43 [1]** | NA | NA |
| B | Healthy controls | 90 ± 26 | 91 ± 29 | 95 ± 32 | 90 ± 29 |
| | unmed. MDD | 92 ± 38 | 92 ± 39 | 95 ± 44 | 92 ± 35 |
| | 7-d med. MDD | **105 ± 45 [1] [2]** | **100 ± 49 [1]** | **107 ± 49 [1] [2]** | **105 ± 51 [1] [2]** |
| C | Healthy controls | 88 ± 32 | 86 ± 34 | 90 ± 37 | 85 ± 31 |
| | 7-d med. MDD | **107 ± 48 [1]** | **108 ± 49 [1]** | **105 ± 46 [1]** | **107 ± 44 [1]** |
| | 28-d med. MDD | **106 ± 51 [1]** | **95 ± 41 [1]** | **104 ± 51 [1]** | **105 ± 51 [1]** |

*Mean durations ± SD (min) are presented, MDD – major depressive disorder, unmed. – unmedicated, med. – medicated, [1] – statistically significant p-values of the t-test that compares a given group to age-matched controls, [2] – statistically significant p-values of the t-test comparing medicated and unmedicated states of MDD patients (dataset B). F – frontal (averaged over F3 and F4), C – central (averaged over C3 and C4), P – parietal (averaged over P3 and P4), O – occipital (averaged over O1 and O2) electrodes.*

**Intra-fractal method reliability**

To assess the intra-fractal method reliability, we correlated between the durations of fractal cycles (i.e., the time interval between two adjacent local peaks) calculated as defined in the main text, i.e., using a minimum peak prominence of 0.94 z and smoothing window of 101 thirty-second epochs, with those calculated using a minimum peak prominence ranging from 0.86 to 1.20 z with a step size of 0.04 z and smoothing windows ranging from 81 to 121 thirty-second epochs with a step size of 10 epochs (Table S7). We found that fractal cycle durations calculated using adjacent minimum peak prominence (i.e., those that differed by 0.04 z) showed r's > 0.92, while those calculated using adjacent smoothing windows (i.e., those that differed by 10 epochs) showed r's > 0.84.

In addition, we correlated fractal cycle durations defined using different channels and found that the correlation coefficients ranged between 0.66 – 0.67 for all datasets except Dataset 2, which showed lower than expected (while still significant) correlations coefficients in the range of 0.42 – 0.45 (Table S1).

Thus, most of the correlations performed to assess intra-fractal method reliability showed correlation coefficients (r > 0.6) higher than those obtained to assess inter-method reliability (r = 0.41 – 0.55), i.e., correlations between fractal and classical cycle durations (Table 2 and Fig. 2 C of the main text and Table S7 here). The strongest correlation between the durations of fractal vs classical cycles (r's > 0.45) were obtained while using the minimum peak prominence of 0.94 and 0.98 z-values, smoothing windows of 101 and 111 (i.e., 50.5 and 55.5 minutes, Table S7) and frontal channels (Table S1). For a discussion on potential sources of differences, see the section "Sources of discrepancy between fractal and classical methods" below.

**Table S7: Intra-fractal method reliability**

| Peak prominence variation | | 0.86 z | 0.90 z | 0.94 z | 0.98 z | 1.20 z |
|---|---|---|---|---|---|---|
| Correlations, r | **0.86 z** | - | 0.93 | 0.86 | 0.82 | 0.55 |
| | **0.90 z** | - | - | 0.92 | 0.88 | 0.60 |
| | **0.94 z** | - | - | - | 0.95 | 0.67 |
| | **0.98 z** | - | - | - | - | 0.74 |
| | **1.20 z** | - | - | - | - | - |
| | Classical cycles | 0.45 | 0.45 | 0.46 | 0.46 | 0.31 |
| Duration, min | Fractal cycles | 88.6 | 90.5 | 92.7 | 93.8 | 106.1 |
| | Classical cycles | 91.0 | | | | |
| **Smoothing window, 30-s epochs** | | **81** | **91** | **101** | **111** | **121** |
| Correlations, r | **81** | - | 0.85 | 0.69 | 0.59 | 0.49 |
| | **91** | - | - | 0.84 | 0.72 | 0.63 |
| | **101** | - | - | - | 0.87 | 0.78 |
| | **111** | - | - | - | - | 0.85 |
| | **121** | - | - | - | - | - |
| | Classical cycles | 0.32 | 0.37 | 0.45 | 0.45 | 0.39 |
| Duration, min | Fractal cycles | 84.5 | 88.6 | 92.4 | 94.5 | 98.6 |

*Fractal cycle durations were calculated using the minimum peak prominence from 0.86 to 1.20 z with the step of 0.04 z and smoothing window from 81 to 121 epochs with the step of 10 thirty-second epochs, r's higher than 0.7 are considered as strong correlation scores, values lower than 0.3 are considered as weak, r's values in the range of 0.3 – 0.7 are considered as moderate scores, all correlations were statistically significant, therefore p-values are not reported, r – Spearman correlation coefficients.*

**Intra-classical method reliability**

To assess the intra-classical method reliability, we correlated between the durations or numbers of classical cycles assessed by two independent human scorers and the automatic sleep cycle detection algorithm, namely, the R "SleepCycles" package (Blume & Cajochen, 2021). The results are presented in Table S8.

In the pooled dataset, the correlation coefficient between classical cycle durations assessed by the two human scorers was 0.8, ranging from 0.7 to 0.9 in different datasets (in literature, r's > 0.7 are interpreted as strong correlations). This is consistent with the literature on sleep staging reporting an average inter-rater agreement of ~ 82.6% (Rosenberg & Van Hout, 2013).

In the pooled dataset, correlation coefficients obtained for sleep cycle durations between human raters and the automatic algorithm showed remarkably lower coefficients of around 0.55 – 0.59, ranging from 0.30 to 0.69 in different datasets ("moderate" correlations). These coefficients were remarkably lower compared to the coefficients obtained between two human scorers ("strong" correlations). The lowest human-automatic inter-rater agreement (r = 0.3) was observed for Dataset 4, where, notably, the data was collected at participants' homes by participants.

In summary, intra-classical method correlations between classical cycle durations scored by two human raters were stronger than the inter-method correlations between fractal and classical cycle durations, which ranged from 0.41 to 0.55 (r's in the range of 0.3 – 0.7 are considered moderate correlations). The human-automatic correlation coefficients for classical cycle durations (r ~ 0.57) lay within the range of the correlation coefficients between fractal and classical cycle durations (r ~ 0.49). In other words, here, the strength of the intra- and inter-method correlations was comparable. For a discussion on potential sources of differences, see the section "Sources of discrepancy between fractal and classical methods" below.

### Table S8: Intra-classical method reliability

| Characteristic | | Scorer | Dataset 1 | Dataset 2 | Dataset 3 | Dataset 4 | Dataset 5 | Pooled dataset |
|---|---|---|---|---|---|---|---|---|
| Classical sleep cycle duration, min | Mean ± SD | Scorer 1 | 87.8 ± 12.6 | 91.5 ± 11.7 | 90.0 ± 13.3 | 93.3 ± 11.0 | 93.8 ± 15.1 | 91.6 ± 13.2 |
| | | Scorer 2 | 88.2 ± 11.6 | 89.4 ± 9.7 | 91.6 ± 11.9 | 90.0 ± 11.9 | 90.4 ± 14.6 | 89.8 ± 12.3 |
| | | Automatic | 94.2 ± 17.7 | 93.0 ± 13.0 | 94.7 ± 10.6 | 101.9 ± 17.9 | 101.2 ± 18.7 | 97.4 ± 16.6 |
| | Correlation, r | Scorer 1–2 | 0.777 | 0.913 | 0.715 | 0.687 | 0.859 | 0.810 |
| | | Scorer 1–automatic | 0.669 | 0.621 | 0.530 | 0.447 | 0.584 | 0.593 |
| | | Scorer 2–automatic | 0.604 | 0.686 | 0.641 | 0.297 | 0.547 | 0.549 |
| No. classical cycles | Mean ± SD | Scorer 1 | 4.4 ± 0.8 | 4.6 ± 0.7 | 4.6 ± 0.6 | 4.7 ± 0.8 | 4.9 ± 0.8 | 4.7 ± 0.8 |
| | | Scorer 2 | 4.2 ± 0.7 | 4.6 ± 0.7 | 4.4 ± 0.6 | 4.7 ± 0.8 | 4.9 ± 0.8 | 4.6 ± 0.8 |
| | | Automatic | 4.1 ± 0.9 | 4.3 ± 0.9 | 4.4 ± 0.6 | 4.2 ± 1.1 | 4.2 ± 1.0 | 4.2 ± 0.9 |
| | Correlation, r | Scorer 1–2 | 0.828 | 0.928 | 0.755 | 1.000 | 0.986 | 0.922 |
| | | Scorer 1–automatic | 0.816 | 0.793 | 0.755 | 0.781 | 0.629 | 0.713 |
| | | Scorer 2–automatic | 0.840 | 0.761 | 0.885 | 0.781 | 0.669 | 0.720 |

*± shows mean and SD, r – Spearman's correlation coefficient, "skipped" cycle – a cycle where a rapid eye movement sleep episode is expected to appear except that it does not.*

### Table S9: Skipped cycle human inter-scorer agreement

| Scorer | Dataset 1 | Dataset 2 | Dataset 3 | Dataset 4 | Dataset 5 | Dataset 6 | Pooled dataset |
|---|---|---|---|---|---|---|---|
| Scorer 1, Number of Skipped Cycles (%) | 5/38 (13%) | 7/39 (18%) | 1/32 (3%) | 19/34 (56%) | 16/62 (26%) | 10/21 (48%) | 58/226 (26%) |
| Scorer 2, Number of Skipped Cycles (%) | 6/38 (15%) | 7/39 (18%) | 3/32 (9%) | 19/34 (59%) | 19/62 (31%) | 10/21 (48%) | 64/226 (28%) |
| Scorer 1 – Scorer 2 Agreement, % | 83 | 100 | 33 | 100 | 84 | 100 | 91 |
